# Single cell CRISPR base editor engineering and transcriptional characterization of cancer mutations

**DOI:** 10.1101/2022.10.31.514258

**Authors:** Heon Seok Kim, Susan M. Grimes, Anuja Sathe, Billy T. Lau, Hanlee P. Ji

## Abstract

We developed a multiplexed single cell technology to genome engineer mutations, directly delineate their genotype among individual cells and determine each mutation’s transcriptional phenotype. This approach uses CRISPR base editors to introduce predesignated variants into a target gene. Long-read sequencing of the target gene’s transcript identifies the engineered mutations among individual cells. Simultaneously, we analyzed the transcriptome profile from the same set of cells by short-read sequencing. By integrating the two types of data, we determined the mutations’ genotype and expression phenotype at single cell resolution. Using cell lines, we engineered and evaluated the phenotype of more than 100 *TP53* mutations. Based on the single cell gene expression, we classified the mutations as having a functionally significant phenotype versus the wild-type state. We validated these results on a subset of mutations using isolated clones analyzed with RNA-seq. Overall, we successfully demonstrated single cell mutation engineering and phenotypic assessment.

## INTRODUCTION

Ongoing genomic studies of cancer are cataloguing extensive numbers of somatic variants. For example, genome sequencing studies have identified numerous cancer mutations across a wide spectrum of tumor types. Many of these mutations result in amino acid substitutions. Given the sheer number of discovered mutations, determining the phenotype of cancer substitutions with functional characterization remains an enormous challenge. In-silico functional predictions of cancer mutations are frequently used as a solution. However, these computational methods do not provide more discrete biological characterization. There remains a significant need for high throughput approaches to functionally evaluate many mutations in an efficient manner. CRISPR base editors and single guide RNAs (**sgRNAs**) have been used for genetic screens - they directly introduce specific variants into target genes at their native genomic loci among transduced cells.^1-4^ These studies examined the altered cellular fitness resulting from the introduced genetic variants, either by counting sgRNA or barcode sequences among the cell pool. However, these approaches do not directly verify the presence of an engineered mutation, since the association with a genotype is imputed based on the sgRNA or the barcode sequence.

Base editors can introduce multiple variants into a target genomic sequence. Although a given sgRNA sequence is intended to generate a single variant, the actual base editing process introduces multiple different, unintended variants at the target genomic sequence. For example, when using the cytosine base editor (**CBE**), the conversion of the either a C to T or a C to G produces different variants other than what was intended. CBEs exhibit cytosine editing in both the target and neighboring bystander cytosines in the editing window with the outcome being multiple different variants at the target sequence site. This variability points to the need to directly genotype the base editor target site as the best approach for verifying the intended mutation being present. Direct validation of an engineered mutation is a necessary step if one is to accurately determine the phenotype. This type of analysis requires examining individual cells.

Several studies have employed a reporter system to infer the presence of the engineered mutations.^3, 4^ However, this method is an indirect approach and assumes the same genome edit has occurred in reporter and endogenous site. Also, these methods may not reflect the precise effects of mutations on gene expression. Citing another example, the single cell Perturb-seq method was adapted to exogenously express genes in the form of cDNAs, containing a specific variants and indirectly measure the mutated gene using barcode sequences.^5^ Although they interrogated the resultant single cell transcriptome changes induced by each variant, this approach had limitations. Specifically, the gene variant was expressed with an exogenous promoter which is not under canonical genetic regulation at the gene’s native locus. Second, variants were delivered to cells with wild-type gene expression of the target gene – this can mask the effect of the variant on protein function. Third, only the barcode sequence is detected instead of the variant itself. Template switching in lentivirus packaging can induce swapping of the variant-barcode association.^6^ This can lead to artifacts in identification and transcriptional phenotyping.

We developed a new method that addresses these challenges and resolves these issues. This method is referred to as transcript-informed single-cell CRISPR sequencing (**TISCC-seq**). This approach relies on CRISPR base-editors to introduce multiple endogenous genetic variants into a given genomic target. Long-read sequencing identifies these mutations directly from a target’s transcript sequence at single cell resolution. Then, we integrate the short-read transcriptome profile from the same single cells (**Figure 1A**). This integrative approach enables single cell direct genotyping and phenotyping of various genetic variants introduced into the native gene locus. Single cell characterization allows one to distinguish the base editor’s intended versus unintended mutations among individual cells We applied this approach to engineer in a series of previously reported cancer mutations in *TP53*, the majority never having been functionally characterized.

**Figure 1.**
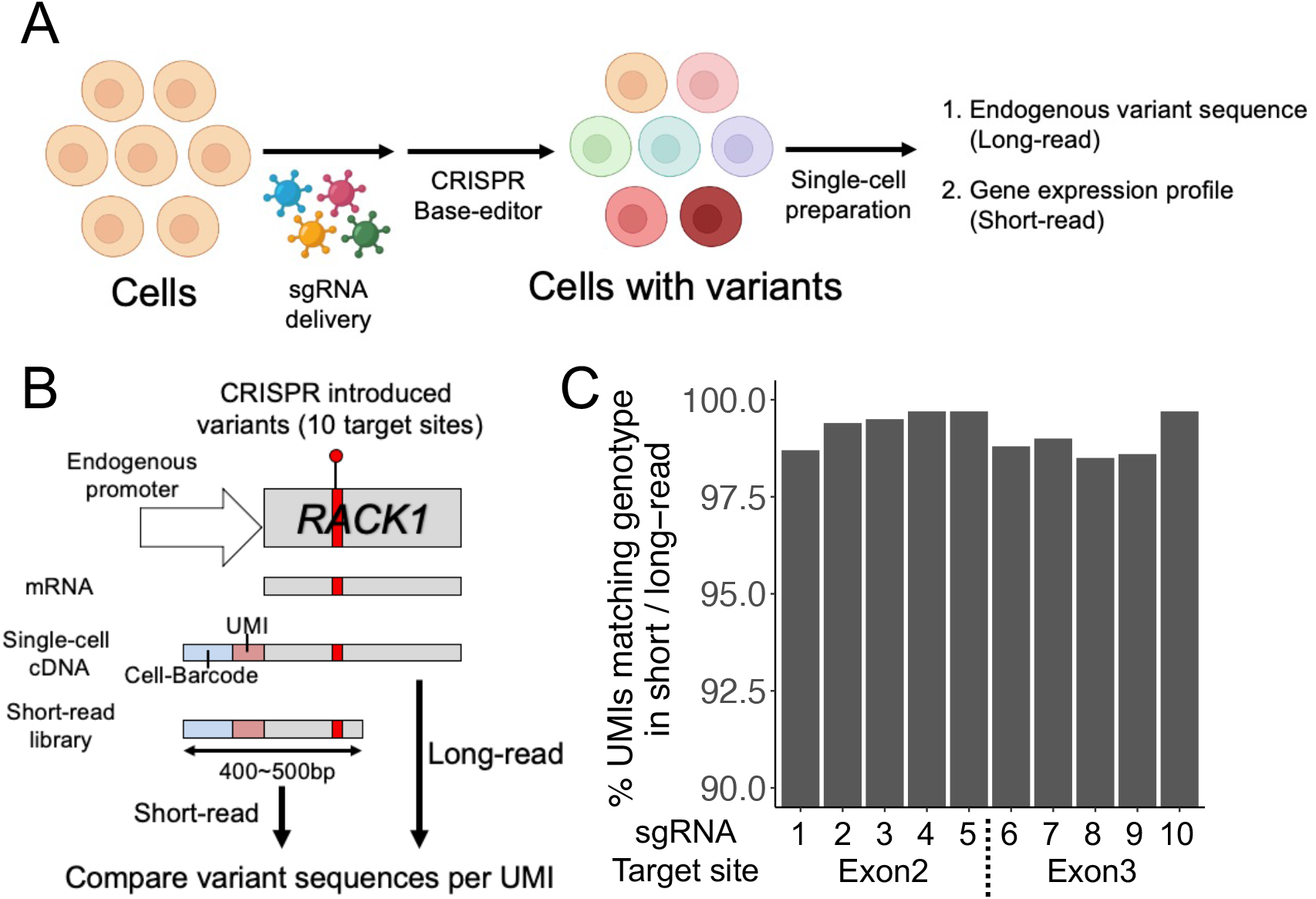
(a) Overview of direct detection and phenotyping of various *TP53* coding mutations. (b) Schematic of the variant calling accuracy comparison between short- and long-read single-cell sequencing. (c) Accuracy of the mutation calling of long-read sequencing. We compared mutation sequences of each sgRNA target site, and calculated proportion of UMIs which have same sequence in short- and long-read sequencing.

## RESULTS

### Identifying CRISPR-induced variants with single cell, long read cDNA sequencing

We conducted an analysis comparing long versus short read single cell cDNA sequencing. For this initial test, we designed an assay to introduce different genetic variants in exon2 and 3 of the *RACK1* gene (**Figure 1B**). The length of *RACK1* cDNA up to exon3 is approximately 500bp – this length interval can be fully covered with short reads. This gene is one the most highly expressed in the HEK293T cell line as determined from single-cell short- and long-read gene expression data from our previous publication.^7^ We designed 10 sgRNAs targeting exon2 and 3 of *RACK1* gene and transduced lentiviruses encoding those sgRNAs to HEK293T cells at 0.1 multiplicity of infection (**Supplementary Table 2**). Transduced cells were selected by puromycin. Then, we transfected a plasmid encoding an adenine base editor (**ABE**) into the cells. This step introduced multiple genetic variants at sgRNA target sites. After six days, we generated single cell cDNAs and extracted genomic DNA from cells derived from the same suspension.

From the genomic DNA of transduced cells, we amplified exon2 or 3 of the *RACK1* gene and performed short-read sequencing to evaluate the frequency of genetic variants in *RACK1* genomic DNA. Based on the DNA sequencing, we identified genetic variants introduced by all of the ten sgRNAs. The frequency of ABE-induced genetic variants varied from 1.1% to 10.1% from the genomic DNA of pooled cells (**Supplementary Figure 1**).

Next, we evaluated the presence of these new variants at single-cell transcript level using single cell cDNAs. These engineered variants were proximal to the 5’ end of the cDNA, allowing us to sequence them with short reads (i.e., Illumina). Short read sequences have a high base quality for variant calling and allowed us to compare the long and short read results. From the single-cell cDNA library, we prepared sequencing libraries for both short- and long-read sequencing to assess single-cell level genetic variants from the *RACK1* transcripts. For short-read sequencing, we amplified exon2 or 3 of *RACK1* from single cell cDNA with cell barcodes and unique molecular index (**UMI**) sequences using the 5’ adaptor primer and exon specific primers (**Figure 1B**). These libraries were sequenced on the Illumina Miseq platform. Similar to regular single-cell gene expression sequencing, we used 26bp of read1 sequences for cell barcode and UMI extraction. The read2 sequences were used for the evaluation of the newly introduced *RACK1* genetic variants at target sites. Using the genetic coordinates of the sgRNA target window (i.e., 3bp to 8bp), for a given read, we identified the corresponding cell barcode, UMI and the genetic variant.

For long-read sequencing, we amplified the entire *RACK1* cDNA using the 5’ adaptor and primers specific to the last 3’ exon from the same single cell cDNA library (**Figure 1B**). The intact cDNA amplicon was sequenced with an Oxford Nanopore instrument. Guppy was used for base calling and minimap2 was used for alignment.^8, 9^ Each sequence read had the cell barcode, UMI and complete *RACK1* cDNA sequence. We extracted the cell barcodes and UMI as we previously described.^7^ After genome alignment of the long-read data, the cell barcodes and UMI fell into soft-clipped sequence. Therefore, we extracted the soft-clipped portion of each read and compared that with the cell barcodes identified from gene expression library sequencing. Only reads with perfectly matching cell barcodes were used for further analysis. Using the aligned long-read data, we identified the *RACK1* genetic variants. Therefore, long read information provided the genetic variants with accompanying cell barcode and UMI sequence. For additional quality control filtering, UMIs with less than three reads were filtered out. We generated consensus genetic variants for each UMI using multiple reads.

We compared the *RACK1* variant calls from short- and long-read single cell data. We analyzed consensus *RACK1* genetic variants for each cell barcode and UMI combination. Across all target sites, we compared 479,509 UMIs: 99.2% of them had identical genetic variants in average (**Figure 1C)**. This result demonstrated the high accuracy of long read identification of CRISPR-engineered genetic variants. Recent improvements in the accuracy of nanopore sequencing and UMI based consensus generation enabled this analysis. We then compared the frequency of genetic variants from genomic DNA and aggregated single-cell cDNA for each of the 10 target sites introduced by base editors. The frequency of each variant between genomic DNA and single-cell cDNA had a high correlation (R^2^ = 0.63, **Supplementary Figure 1**).

### Base editor guide RNA designs for *TP53* cancer mutations

We introduced a set of sgRNAs designed for multiple *TP53* mutations and used TISCC-Seq to obtain the gene expression profile and *TP53* genotype from individual cells. First, we focused on the design of the genome engineering of *TP53* mutations (**Figure 2A**). We identified *TP53* mutations which were reported more than nine times in the COSMIC database.^10^ The majority of these frequent cancer mutations were within the *TP53* DNA-binding domain. The total number of coding mutations was 351. We designed base editor libraries targeting this mutation set. To cover as many mutations as possible we used several base editor combinations: (1) CBE with NGG protospacer adjacent motif (**PAM**); (2) CBE with a NG PAM; (3) ABE with NGG PAM; (4) ABE with a NG PAM. Using the NGG PAM base editors, we designed 74 sgRNAs targeting 99 *TP53* variants. The NG PAM base editors have more flexible PAM, so we were able to design an additional 88 sgRNAs targeting 159 variants (**Supplementary Figure 2**). Most of sgRNAs targeted the DNA binding domain of TP53 protein (**Figure 2B**).

**Figure 2.**
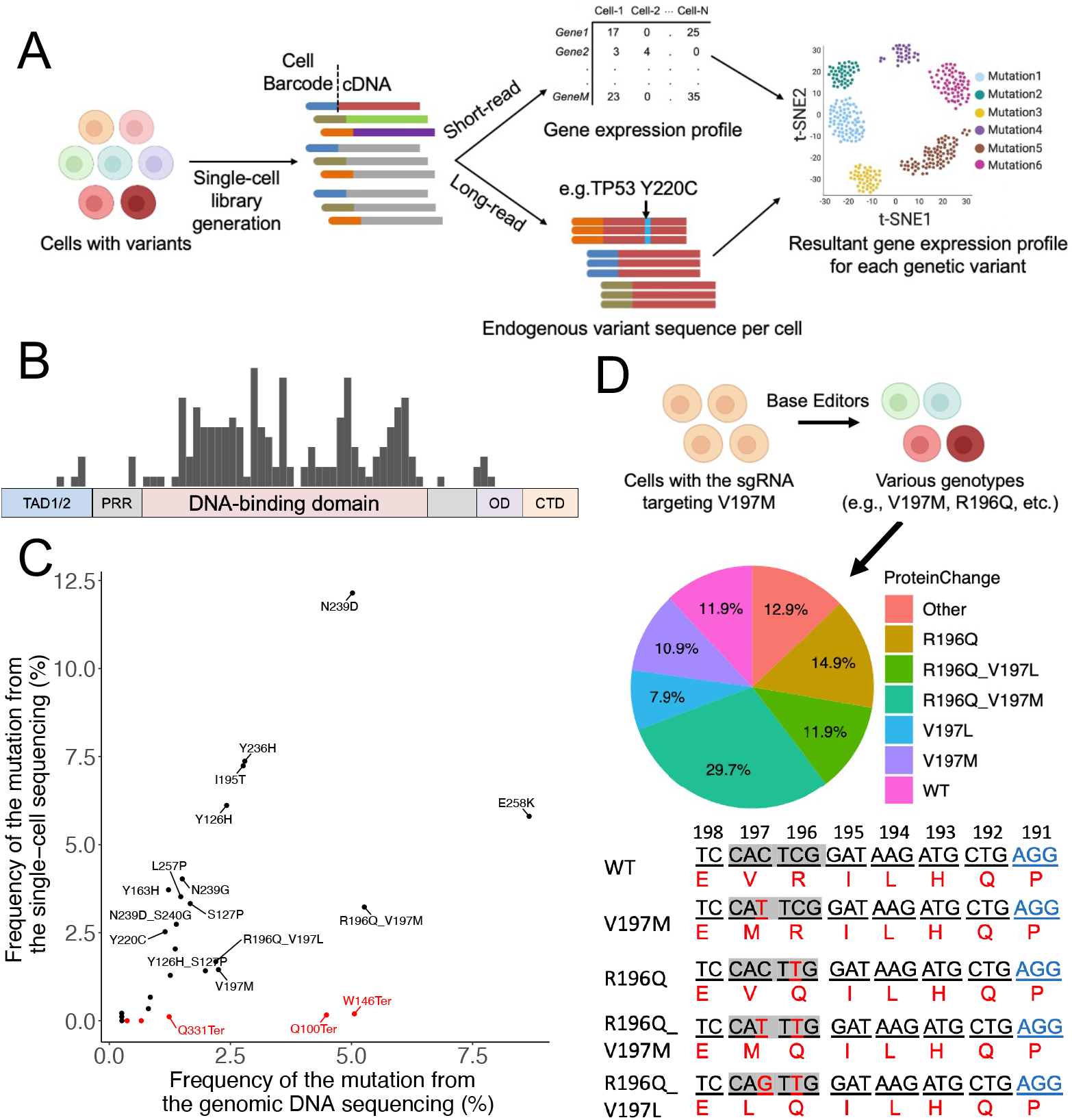
(a) Overview of single-cell cDNA analysis pipe-line. (b) Structure of P53 protein and distribution of sgRNA target sites we used in this study. TAD, transactivation domain; PRR, proline-rich region; OD, oligomerization domain; CTD, carboxyl terminus domain. (c) Dot plot showing the proportion of each genetic variant detected from single-cell cDNA and genomic DNA. Red dots represent variant with premature stop codon. (d) Cells with same sgRNA can result in various genotypes. The pie chart shows the proportion of resultant amino acid changes from cells with sgRNA targeting V197M mutation. Proportions of mutations are calculated from the single-cell cDNA long-read sequencing. Underlines indicate each triplet codon and number indicate position of the codon. Red DNA sequences indicate substituted bases and blues indicate PAM sequences.

Base editors can alter any target nucleotide in their target window (i.e., 3bp to 8bp) which leads to different nucleotides at that position. TISCC-seq identified this variation among single cells. For example, the sgRNA introducing E258K mutation by C to T substitution induces the E258G mutation by C to G substitution (**Supplementary Figure 3**). Similarly, the sgRNA introducing S127P mutation by A to G substitution at the 3^rd^ adenine induces the Y126H mutation by A to G substitution at the 6^th^ adenine (**Supplementary Figure 3**). Therefore, this result suggests that any given sgRNA can introduce multiple variants depending on the window sequence context. The entire number of amino acid changes that could be introduced by the NGG or NG PAM base editors and our sgRNA libraries were 920 and 1999 respectively. For the final design, we targeted 251 known *TP53* mutations with the potential for introducing 2892 possible amino acid changes (**Supplementary Figure 2**).

### CRISPR base editor engineering of *TP53* mutations

We used HCT116 and U2OS human cell lines for this study. Both cell lines have wild-type *TP53* which we independently confirmed.^11-13^ The P53 pathway is repressed by the negative regulator MDM2 in both cell lines.^14^ The oncoprotein MDM2 is an E2 ubiquitin ligase.^15^ It binds to and promotes the ubiquitin-dependent degradation of the TP53 protein. The small molecule nutlin-3a can inhibit P53-MDM2 binding efficiently.^16^ Therefore, we used nutlin-3a to activate the P53 pathway and select for *TP53* mutations.

We generated four sgRNA libraries for each base editor (NGG-CBE, NGG-ABE, NG-CBE, NG-ABE) – the combined libraries were designed to cover the preselected *TP53* mutations. We transduced those libraries using a lentivirus system to both the HCT116 and U2OS cell lines. The cells were transfected with each respective base editor plasmids. It had been reported that base editors can induce off-target RNA editing.^17^ To minimize those effects, we chose transient transfection rather than stable expression of base editors. Typically, plasmid based protein expression peaks after 24hrs of transfection and diminishes after 5-6 days.^18^ Six days after transfection, we used nutlin-3a to activate the TP53 pathway.

### TISCC-seq detection of *TP53* mutations

After 10 days of nutlin-3a treatment, we harvested the cells for suspension, prepared single-cell cDNA libraries and also extracted genomic DNA from a portion of the cell suspension **(Methods**). We amplified *TP53* transcripts from the single-cell cDNA library, sequenced their full-length transcript and determined the presence of the *TP53* mutation from the long read data (**Figure 2A**). As an important additional step, we extracted cell barcodes and UMI per each long-read as described earlier. To prevent the effect of sequencing error in UMI region, we filtered out any UMI with less than 10 long reads. As a quality control threshold, we used only the cell barcode and UMI combinations found in 10 or more reads. For generating a consensus, we also included UMIs with a low edit distance, assuming the differences were related to sequencing errors. For *TP53* variant calling, we extracted every nucleotide sequence in the sgRNA target window (e.g., chr 17:7674940-7674945 for the sgRNA in **Figure 2D**) and compared them with reference sequence (e.g., CACTCG to CATTCG). Based on nucleotide changes of a given mutation, we determined the amino acid substitution at the target site (e.g., V196M).

For independent validation, we used amplicon sequencing from the transduced cells’ genomic DNA to independently assess the frequency of a subset of *TP53* mutations. This analysis compared the frequency of each *TP53* mutations introduced by 12 sgRNAs in genomic DNA versus the results from analyzing the single-cell cDNA from HCT116 cells. These *TP53* mutations were introduced efficiently with up to 12.1% for one variant and 27 variants were introduced with a frequency greater than 0.25%. The prevalence of each mutation from single-cell cDNA and genomic DNA was generally correlated (**Figure 2C, Supplementary Figure 4**, R^2^ = 0.59). Some variants had higher frequency in genomic DNA and lower in cDNA (i.e., W146Ter). This result means that for some mutations the corresponding transcripts were not expressed efficiently or were subjected to higher RNA degradation. The lower prevalence of cDNA mutations may reflect effects from nonsense mediated decay (**NMD**). This process is a surveillance mechanism that eliminates mRNA transcripts containing premature stop codons. For example, although 5.1% of cells had a W146Ter mutation at the genomic DNA level, this mutation was not detected as frequently at the single cDNA level (0.2%) because the transcripts with the variant were degraded in cells by NMD (**Figure 2C**).

As another type of validation, we also sequenced the sgRNA expressed in each cell from single-cell cDNA using a direct capture method previously described.^7, 19^ Most of the single-cell CRISPR screen studies have relied on an sgRNA sequencing method to infer the resultant genetic edits.^20-23^ This method assumes that cells with the sgRNA have the targeted genomic edit. However, the efficiency of base editors is lower than Cas9 nuclease.^24, 25^ As described earlier, a base editor may introduce multiple genetic variants from the same sgRNA (**Supplementary Figure 3**). Therefore, one cannot assume that cells transduced with base editors and a single sgRNA have the intended variant at the target position (**Figure 2D and Supplementary Figure 5**). Our results showed that this was the case. For example, we evaluated a sgRNA which was designed to introduce the *TP53* V197M mutation. The sgRNA’s target site has three cytosines in its window. Among 101 cells expressing this specific sgRNA, 11 cells had V197M mutation while 30 cells had both R196Q and V197M mutations (**Figure 2D**). Therefore, the conventional single-cell CRISPR screening method using sgRNA sequencing did not correctly identify the introduced variants among the various single cells. In contrast, with direct long read sequencing of the full-length target transcripts from single cells, we bypassed this issue and directly identified the actual mutation introduced by the base editor from the cDNA.

### TISCC-seq and expression analysis of HCT116 cells with *TP53* mutations

We performed gene expression analysis using the same single-cell cDNA library we used for long-read sequencing. As we described previously, we integrated the single cell *TP53* mutation genotypes from long reads with the single-cell gene expression profile data from short reads.^7^ We used cell barcode matching between the long read data with a mutation genotype and the short read data **(Methods**). This process allowed us to link those cells with *TP53* mutation to their individual gene expression profiles. To conduct a cluster analysis of the cells with different *TP53* mutations, we used Uniform Manifold Approximation and Projection (**UMAP**) (**Figure 3**). We investigated the effect of P53 pathway activation by nutlin-3a in HCT116 cells with *TP53* mutations using a subset of our sgRNA library (10 sgRNAs). When we compared the gene expression profiles between cells with wild-type or *TP53* mutations, there was a significant and clearly delineated difference upon P53 pathway activation (**Figure 3A and 3B**). Overall, cells with deleterious *TP53* mutations had lower *TP53* related gene expression compared to the wild-type cells (**Supplementary Figure 6**).

**Figure 3.**
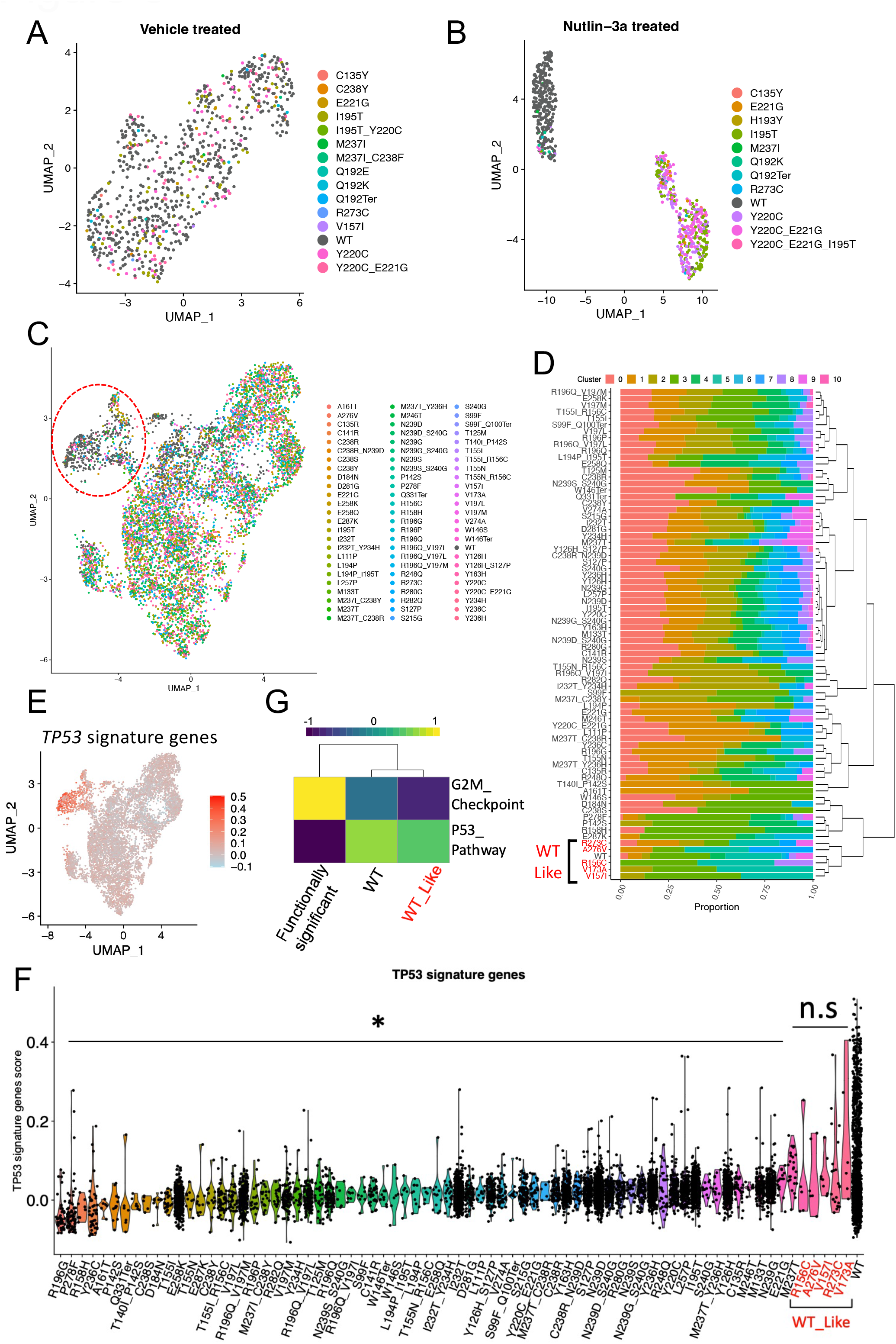
(a, b, c) UMAP plot showing single-cell gene expression profile per each genetic variant. HCT116 cells are treated with vehicle (a) or Nutlin-3a (b) after the introduction of variants using subset of sgRNA library. (c) HCT116 cells are treated with Nutlin-3a after introduction of variants using full sgRNA library. (d) Proportion of UMAP cluster from cells with each genetic variant. Hierarchical clustering was performed based on the proportion to categorize genetic variants. Reds indicate wild type-like variants. (e) UMAP embedding of cells colored by P53 pathway gene scores. (f) Violin plot showing P53 pathway gene score per cells with each genetic variant. *: P < 0.03, n.s: Not significant; two-sided t-test. (g) Heatmap showing average GSVA enrichment score of selected Hallmark pathways per each category of genetic variant.

Next, we sequenced HCT116 cells transduced with our full *TP53* sgRNA library and activated by nutlin-3a. Among the 42,564 cells that were sequenced, we generated high quality long read UMIs (UMI read count > 9) covering *TP53* from 12,887 cells. For each cell, we had an average of 898 *TP53* reads with 4.5 UMIs. We filtered out cells which had a heterozygous mutation. Overall, we detected total of 169 different mutations distributed among the various single cells.

We analyzed the single cell gene expression for each mutation. To provide a robust measurement of single cell expression, we filtered out those *TP53* mutations expressed in fewer than five cells. This step retained 74 mutations for further analysis. As is clear via UMAP clustering, the cells with wild-type versus *TP53* mutations clearly separated among different clusters. Wild-type cells were predominantly clustered in Cluster 5 and 9 (**Figure 3C and Supplementary Figure 7**). For each variant, we calculated its proportion within each cluster and performed hierarchical clustering of each variant based on the proportion (**Figure 3D**). Cells with the following five mutations (R156C, V157I, V173A, R273C and A276V) clustered with the wild type cells. This result was a preliminary indication that this set of mutations did not have a significant impact on the gene expression phenotype - we annotated them as wild-type like and the others as functionally significant.

We analyzed the expression of 343 genes involved in the P53 pathway across the single cell data (**Supplementary Table 6**).^26^ Cells that were wild type or with mutations that were wild type like had higher expression of P53 pathway signature genes (**Figure 3E**). Wild-type cells had higher P53 pathway gene expressions core compared to the majority of cells expressing functionally significant *TP53* mutations (**Figure 3F**, P < 0.03). Additionally, we analyzed the expression of the *CDKN1A* gene, otherwise generally referred to the p21 protein. CDKN1A is a regulator of cell cycle progression and arrest. Wild-type cells had higher *CDKN1A* expression compared to the cells with functionally significant *TP53* mutations (**Supplementary Figure 8**). Next, we performed pathway analysis between wild-type cell and cells with wild-type like versus functionally significant variants. Cells with functionally significant mutations had lower P53 pathway activity and higher G2M checkpoint gene expression than the wild-type cells (**Figure 3G**, P= 1.66e-11 and 1.66e-11). In addition, cells with wild-type like variants expressing the R156C, V157I, V173A, R273C or A276V did not have differences on two pathways compared to cells with wild type *TP53* (**Figure 3G**, P= 0.95 and 0.44). These results are evidence that this subset of the mutations had features similar to wild type and thus had less functional impact. In summary, wild-type cells had higher active P53 pathway activity and related gene expression than cells with functionally significant *TP53* variants. These results validated the TISCC-seq method for high throughput functional classification of these mutations.

### TISCC-seq analysis of *TP53* mutations in U2OS cell line

As an additional verification of our results, we performed a similar analysis with the U2OS cell line using the same sgRNAs for the *TP53* mutations. Among 38,451 cells that we sequenced, we were able to acquire high quality long-read sequences from 12,155 cells. On average per each cell, we sequenced 890 *TP53* reads with 4.6 UMIs. As described, we applied a filtering strategy to eliminated heterozygous mutations. For the U2OS line, we characterized 161 mutations with TISCC-seq. For gene expression analysis, we used the 62 variants which were detected in more than five cells. From the UMAP analysis, wild-type cells and cells with *TP53* mutations separate into distinct clusters (**Figure 4A** and **Supplementary Figure 9**). Wild-type cells were primarily associated with Cluster 1. For each mutation, we calculated its proportion within each cluster and performed hierarchical clustering based on this cluster proportion (**Figure 4B**). From the hierarchical clustering results, we identified four mutations, T140I, R156C, T221I and R273C, that were associated with wild type *TP53*. The R156C and R273C mutations had a similar association with the wild type cells for both the HCT116 and U2OS cell lines. The wild-type U2OS cells had higher expression of *CDKN1A* and other P53 pathway signature genes compared to the majority of cells expressing functionally significant *TP53* mutations (**Figure 4C, 4D and Supplementary Figure 10**). The analysis of pathway activity showed that cells with functionally significant mutations had significantly lower P53 pathway activity and higher G2M checkpoint gene expression (**Figure 4E**, P= 1.62e-12 and 1.62e-12). Conversely, cells with wild-type like mutations were not statistically significant to the same extreme degree as the functionally significant mutations (**Figure 4E**, P= 0.52 and 0.001).

**Figure 4.**
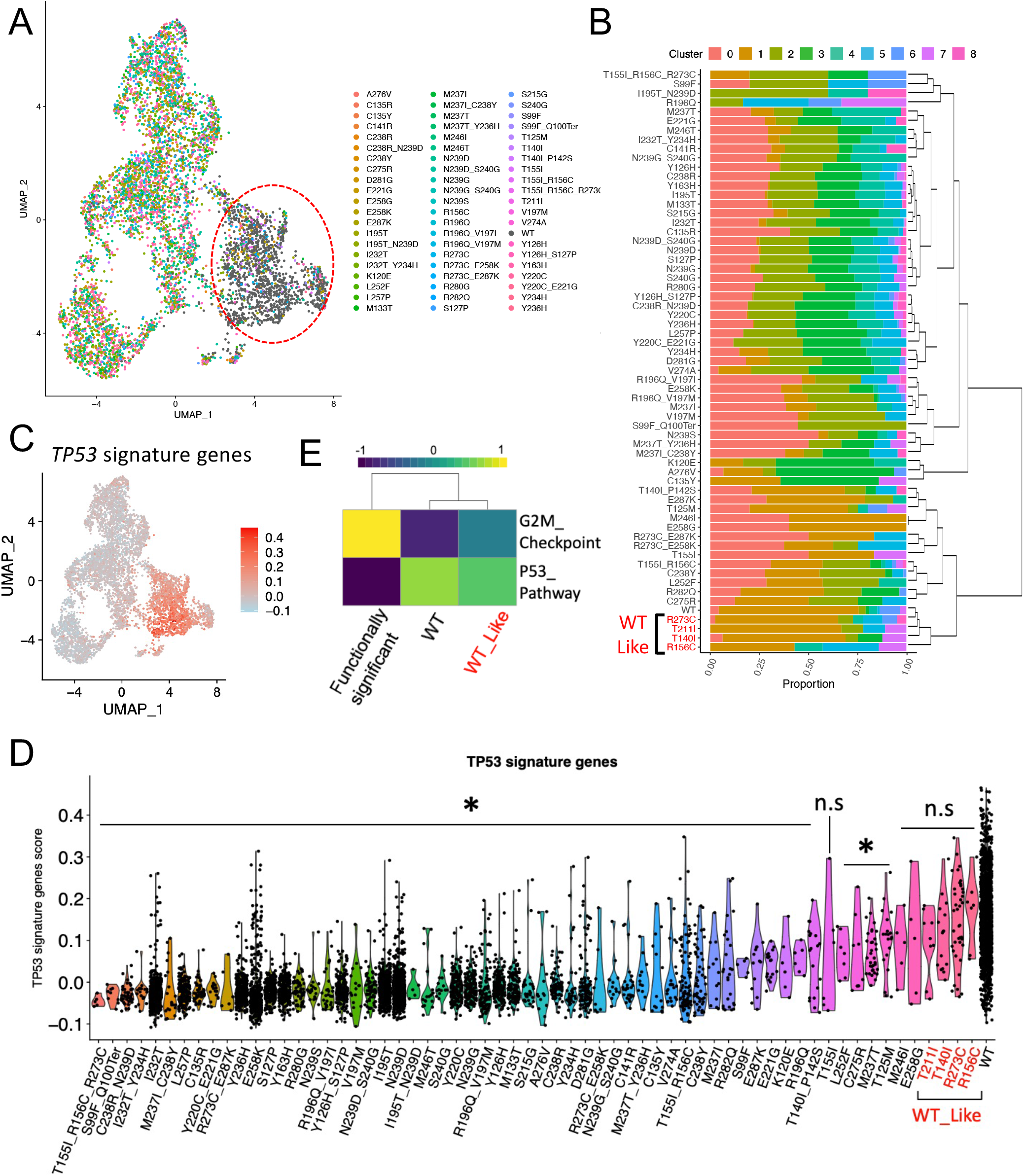
(a) UMAP plot showing single-cell gene expression profile per each genetic variant. U2OS cells are treated with Nutlin-3a after introduction of variants using full sgRNA library. (b) Proportion of UMAP cluster from cells with each genetic variant. Hierarchical clustering was performed based on the proportion to categorize genetic variants. Reds indicate wild type-like variants. (c) UMAP embedding of cells colored by P53 pathway gene scores. (d) Violin plot showing the P53 pathway gene score per cells with each genetic variant. *: P < 0.03, n.s: Not significant; two-sided t-test. (e) Heatmap showing average GSVA enrichment score of selected Hallmark pathways per each category of genetic variant.

### Confirmation of single-cell mutation characterization using clonal cell lines

Our prior experiments were highly multiplexed in engineering different mutations. Providing additional confirmation of the single cell results, we conducted simplex experiments of individual mutations using the HCT116 cell line. Using the ABE, we generated homozygous clonal cell lines with either the *TP53* I195T or Y220C mutation which were functionally significant and had enough cells from single-cell assay. To obtain clones, we used limiting dilution after ABE transfection. These two mutations have been reported to have a deleterious effect on function^10, 27^ and the multiplexed TISCC-seq results also demonstrated that they had a functional effect (**Figure 3D**). We performed bulk-RNA seq from nutlin-3a treated wild-type cells and those clonal cells. We compared the result with single cell results from HCT116 cell-lines (**Figure 5**).

**Figure 5.**
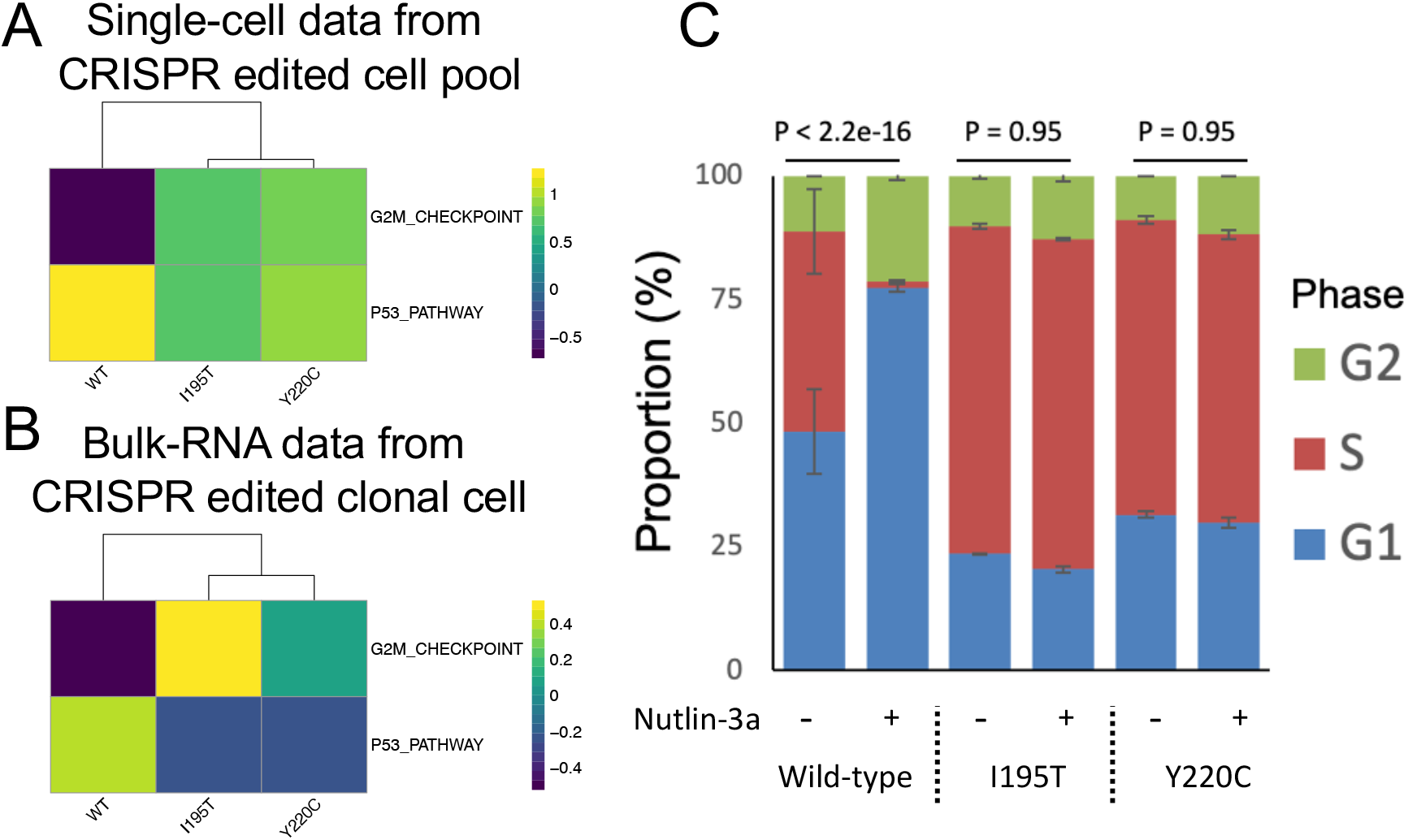
(a, b) Heatmap showing the average GSVA enrichment score of selected Hallmark pathways. (a) Scores are calculated from single-cell analysis of heterogenous *TP53* genetic variants pool. (b) Scores are calculated from bulk RNA sequencing from clonal cells with indicated *TP53* genetic variants. (c) Cell cycle analysis using DNA content staining using clonal cells. Genetic variant per cells and nutlin-3a treatments are indicated (n = 2, mean ± s.e.m.). P values are calculated by Chi-squared test.

From the single cell results, both mutations demonstrated lower P53 pathway activity and higher G2M checkpoint gene expression than wild-type cells (**Figure 5A**, I195T: P = 2.2e-11 and 1.7e-3. Y220C: 2.2e-11 and 9.4e-2). From the conventional, bulk-based RNA-seq results, we observed the same effect on the same pathways (**Figure 5B**, I195T: P = 3.4e-6 and 2.4e-7. Y220C: 1.0e-4 and 2.6e-7). Next, we performed differentially gene expression (**DGE**) analysis between wild-type versus mutation-bearing cells. We compared the DGE results from scRNA-seq and standard RNA-seq. For the I195T or the Y220C mutations, we identified the top 100 genes determined from single cell RNA-seq data. For the I195T mutation, 94 out of 100 were confirmed as showing differentially expression per the conventional RNA-seq. Likewise for the Y220C mutation, 80 out of 100 genes were confirmed as showing differentially expression per the conventional RNA-seq (**Supplementary Figure 11, Supplementary Table 7 and 8**, P < 1.0e-5).

Overall, the I195T and Y220C cell lines had higher G2M checkpoint gene expression as an indicator of more active cell division compared to the cells with wild type *TP53*. To validate this result, we evaluated cell division and cell cycling from wild-type and *TP53*-mutated HCT116 cells using 5-ethynyl-2′-deoxyuridine (**Edu**) and a propidium iodide (**PI**) flow cytometry assay. The PI assay detects total DNA amounts for G1 and G2-phase comparison. The EdU assay labels newly synthesized DNA to detect S-phase. The cell cycle of wild-type HCT116 cells was arrested by nutlin-3a treatment (**Figure 5C and Supplementary Figure 11**, P < 2.2e-16). In contrast, the cell cycle of HCT116 cells with either the I195T or the Y220C mutations did not undergo arrest with nutlin-3a treatment (**Figure 5C and Supplementary Figure 11**, P= 0.95 and 0.95). This result established that this single cell approach accurately identified the phenotypes of these mutations.

## DISCUSSION

In this study, we report a multiplexed method that uses base editors to introduce specific cancer mutations and single cell sequencing to identify the genotype and phenotypes of the induced cancer mutations. Referred to as TISCC-seq, this approach overcomes issues with short-read based single-cell or bulk CRISPR screens, neither of which verify endogenous DNA variants that are engineered into the genomes of cells. This approach integrated single cell long-read and short-read sequencing for CRISPR base editor screens. As a result, endogenous genetic variants introduced by the CRISPR base editor are directly confirmed from the target gene transcript. At single-cell resolution, the genetic variant and its resultant transcriptome changes become evident. Therefore, we can determine the functional consequences of *TP53* mutations across different cell lines. Some mutations had a greater functional impact on the cells’ gene expression while a smaller subset had a wild-type like phenotype. Our results corroborated some in silico predictions. For example, the R156C mutation is predicted to have neutral effect on P53 pathway.^10, 28^ This was confirmed experimentally among our results. In both cell lines used in this study, this mutation had a wild type phenotype. Overall, this approach has the potential for enabling highly multiplexed functional evaluation of cancer mutations and germline variants. Following functional assays using cell-lines with desired genetic variants will help deeper understanding of the phenotype of each variant as shown in **Figure 5**.

Although we used four base editors for this study, there were some mutations that we were unable to target (**Supplementary Figure 2)**. We anticipate that modification of base editor properties such as their enzymatic activity^29, 30^, window^31^ and PAM restriction^32^ will broaden the types of mutations and other variants which can be engineered into genomes. The prime editor which can introduce any genetic variant at the target site will even enable saturation mutagenesis of the target gene.^33^

The complexities of single cell sequencing and its higher cost limit the scalability of single-cell CRISPR screens compared to conventional genetic screens done with conventional bulk assays. TISCC-seq provides some potential benefits that may be useful for standard CRISPR screens. For example, one can use a bulk-based cellular genetic screen for hundreds of thousands sgRNAs generating variants and then narrow down the sgRNAs to the hundreds with significant impact on cell survival or drug response. Then, TISCC-seq can be used to deeper analysis of sgRNAs by detecting genuine endogenous mutations and their resultant phenotype at single-cell level resolution. This combination may enable more accurate evaluation of CRISPR-based screens in the future.

## METHODS

### Cell culture conditions

HEK293T (ATCC CRL-11268) cells were maintained in Dulbecco’s modified Eagle’s medium (DMEM) with 10% fetal bovine serum (FBS). HCT116 (ATCC CCL-247) cells and U2OS (ATCC HTB-96) were maintained in McCoy’s 5A modified medium supplemented with 10% FBS. We stimulated P53 pathway of cells with 10μM of Nutlin-3a. Cells were authenticated by STR profiling. All cell lines were confirmed by PCR to be free of mycoplasma contamination.

### Lentiviral gRNA library production

The oligonucleotides for sgRNA library generation were ordered using IDT oPools Oligo Pools (Coralville, Iowa, USA). Amplified gRNA cassettes were cloned using NEBuilder HiFi DNA Assembly Master Mix (New England Biolabs, Ipswich, MA, USA) into lentiGuide-Puro (Addgene plasmid #52963). Purified plasmids were electroporated to ElectroMAX Stbl4 Competent Cells (New England Biolabs) and amplified.

### Lentivirus production

Approximately 2.0 × 10^6^ HEK293T cells were plated 24h prior to transfection. Cells were transfected with pMD2.G (500 ng, Addgene plasmid #12259), psPAX2 (1500 ng, Addgene plasmid #12260) and lentiviral sgRNA library (2000 ng) using Lipofectamine 2000 (Invitrogen) as per the manufacturer’s protocol. The viral supernatant was collected after 48hr of transfection. The supernatants were filtered through a 0.45μm filter and transduced to cells.

### Lentivirus transduction

HCT116 and U2OS cells were diluted to 1.4 × 10^5^ and 0.7 × 10^5^ cells / mL and plated a day prior to the transduction. Lentiviral supernatant and polybrene (8 μg / mL, Sigma-Aldrich, MO, USA) were added to the cells. After 24 hours, transduced cells were selected by puromycin (Life technologies, CA, USA) at concentration of 0.4 μg / mL and 1.0 μg / mL.

### Transfection and electroporation condition

We used 1.2 × 10^6^ HEK293T cells to transfect the base editor plasmids (2000 ng) using Lipofectamine 2000 (Invitrogen, Carlsbad, CA, USA) as per the manufacturer’s protocol. We used 1.0 × 10^6^ HCT116 or U2OS cells to transfect the base editor plasmids (2600 ng) using SE solution and 4D-nucleofector (Lonza, Switzerland) as per the manufacturer’s protocol. Base editor plasmids pCMV_AncBE4max_P2A_GFP and pCMV_ABEmax_P2A_GFP were gifts from David Liu (Addgene plasmid # 112100 and 112101).^34^ Base editor constructs pCAG-CBE4max-SpG-P2A-EGFP (RTW4552) and pCMV-T7-ABEmax(7.10)-SpG-P2A-EGFP (RTW4562) were gifts from Benjamin Kleinstiver (Addgene plasmid # 139998 and # 140002).^32^ After six days of electroporation, cells were subjected to chemical treatment or single-cell library preparation. For *TP53* variant clone generation, base editor plasmids (2250 ng) and sgRNA plasmid (750 ng) were electroporated to cells. We conducted single cell subcloning with limiting dilution and confirmed the genotype of the target with PCR amplification and sequencing.

### Single-cell library preparation

Single-cell cDNA and gene expression libraries are generated using Chromium Next GEM Single Cell 5’ Library & Gel Bead Kit v2 (10X Genomics, Pleasanton, CA, USA) according to the manufacturer’s protocol. The cDNA and gene expression libraries are amplified with 16 and 14 cycles of PCR respectively. The quality of gene expression libraries is confirmed using 2% E-Gel (ThermoFisher Scientific, Waltham, MA, USA). We quantified the sequencing libraries using Qubit (Invitrogen) and sequenced on Illumina sequencers (Illumina, San Diego, CA, USA).

### Single-cell sgRNA capture and sequencing

The sgRNA direct capture was performed as previously described.^7, 19^ Briefly, six pmol of sgRNA scaffold binding primer was added to RT master mix. After cDNA amplification, the sgRNA fractions were purified using SPRIselect bead (Beckman Coulter Life Sciences, CA, USA). The library was amplified and sequenced with gene expression library.

### Long-read sequencing

Ten ng of the single-cell full length cDNA were used to amplify transcripts. Primer sequences are shown in **Supplementary Table 5**. We used KAPA HiFi HotStart ReadyMix (Roche, Basel, Switzerland) for amplification. Libraries were prepared with 900fmol of each amplicon for Promethion flow cell FLO-PRO002 (Oxford Nanopore Technologies, Oxford, UK) using Native Barcoding Expansion and Ligation Sequencing Kit (Oxford Nanopore Technologies) according to the manufacturer’s protocol. Libraries were sequenced on a Promethion over 72h.

### Single cell transcript analysis

#### Short read transcripts

Basecalling for 5’ gene expression libraries was performed using cellranger 6.0 (10X Genomics). In preparation for integrated analysis, the transcript count matrices generated by cellranger were processed by Seurat 3.0.2^35^ QC filtering removed cells with fewer than 100 or more than 8000 genes, cells with more than 30% mitochondrial genes and cells predicted to be doublets by DoubletFinder.^36^ Additionally, any genes present in three or fewer cells were removed. Batch effects between each single-cell cDNA generation reaction and base editors were corrected by Harmony.^37^ Cell cycle phase were also corrected by Harmony.

#### Long read variant calling

Basecalling was performed using guppy 5 with super accuracy mode and alignment to the GRCh38 reference genome using minimap2.^8, 9^ Cell barcodes and UMIs are extracted as previously described.^7^ For *TP53* mutation calling, we filtered out UMIs less than 10 reads and consolidated UMIs with high similarity (edit distance less than 3). A custom python script utilizing the pysam module was used to identify reads spanning the sgRNA target windows, and extract the base calls at each position within the window. Base calls were used predict amino acid changes per each cell. Cells with heterozygous amino acid changes were excluded for the gene expression analysis. Output from this script was summarized to provide expected amino acid change per cell barcode.

#### Integration of long and short reads

The variant per cell barcode table were added to the Seurat object metadata as a new column. Cells without high-quality long-read data were filtered. For gene expression analysis, we filtered variants which were detected in less than 5 cells. A hierarchical clustering was done in R using hclust, cutree and dendextend. Biological pathway analysis was performed with the Gene Set Variation Analysis (**GSVA**) tool.^38^

### Cell cycle analysis

We used Click-iT™ Plus EdU Alexa Fluor™ 488 Flow Cytometry Assay Kit (Life technologies) according to manufacturer’s protocol. Briefly, we plated cells a day prior to nutlin-3a or vehicle treatment. After 24 hrs of chemical treatment, cells in S-phase were labeled with 10 mM EdU solution for 2 hrs. FxCycle™ PI/RNase Staining Solution (Life technologies) was used for PI staining. After the staining, cells were analyzed by NovoCyte Quanteon Flow Cytometer Systems (Agilent, Santa Clara, CA, USA).

### RNA sequencing

We used KAPA mRNA HyperPrep Kit (Roche) for mRNA sequencing library preparation according to manufacturer’s protocol. For each cell type, we used triplicate library preparations with 1 μg of total RNA as an input. Libraries were sequenced by NextSeq (Illumina) by 75bp paired-end sequencing. The reads were aligned to the reference genome GRCh38 by a two-pass method with STAR and gene expression level was measured using HT-Seq.^39, 40^ We used DESeq2 for DE analysis.^41^ Biological pathway analysis was performed with the Gene Set Variation Analysis (GSVA) tool.^38^

## Data availability

High-throughput DNA sequencing files are available from the NCBI SRA under BioProject PRJNA880341. The URL is listed below: https://dataview.ncbi.nlm.nih.gov/object/PRJNA880341?reviewer=6roem776siu8nui8jv3oq73caf

## Code availability

Scripts for analysis are available on zenodo (https://zenodo.org/badge/latestdoi/555044610) under the MIT license terms.

## Acknowledgements

Portions of this work was supported by US National Institutes of Health grants R33 CA247700 (HPJ, HSK, BTL), R01HG006137 (HPJ, HSK), and R35HG011292-01 to BTL and HSK. HPJ and HSK also received support from the Clayville Foundation.

## Author contributions

HSK and HPJ were involved in conception and design of the study, development of methodology, acquisition of data, analysis and interpretation of data, and writing of the manuscript. HSK, SMG, AS, BTL and HPJ were involved in analysis and interpretation of data. All authors contributed to the writing of the manuscript. HPJ oversaw all aspects of the study. All author(s) read and approved the final manuscript.

