## Supplementary Figures for "Single cell CRISPR base editor engineering and transcriptional characterization of cancer mutations"

**Institutions:**

[1] Division of Oncology, Department of Medicine, Stanford University School of Medicine, Stanford, CA, United States

[2] Department of Electrical Engineering, Stanford University, Stanford, CA, United States

**Corresponding author:**

Hanlee P. Ji

CCSR 1115, 269 Campus Drive, Stanford, CA-94305, USA

**Supplementary Figure 1. Dot plot showing the proportion of each genetic variant detected from single-cell cDNA and genomic DNA from *RACK1* edited HEK293T cells.**

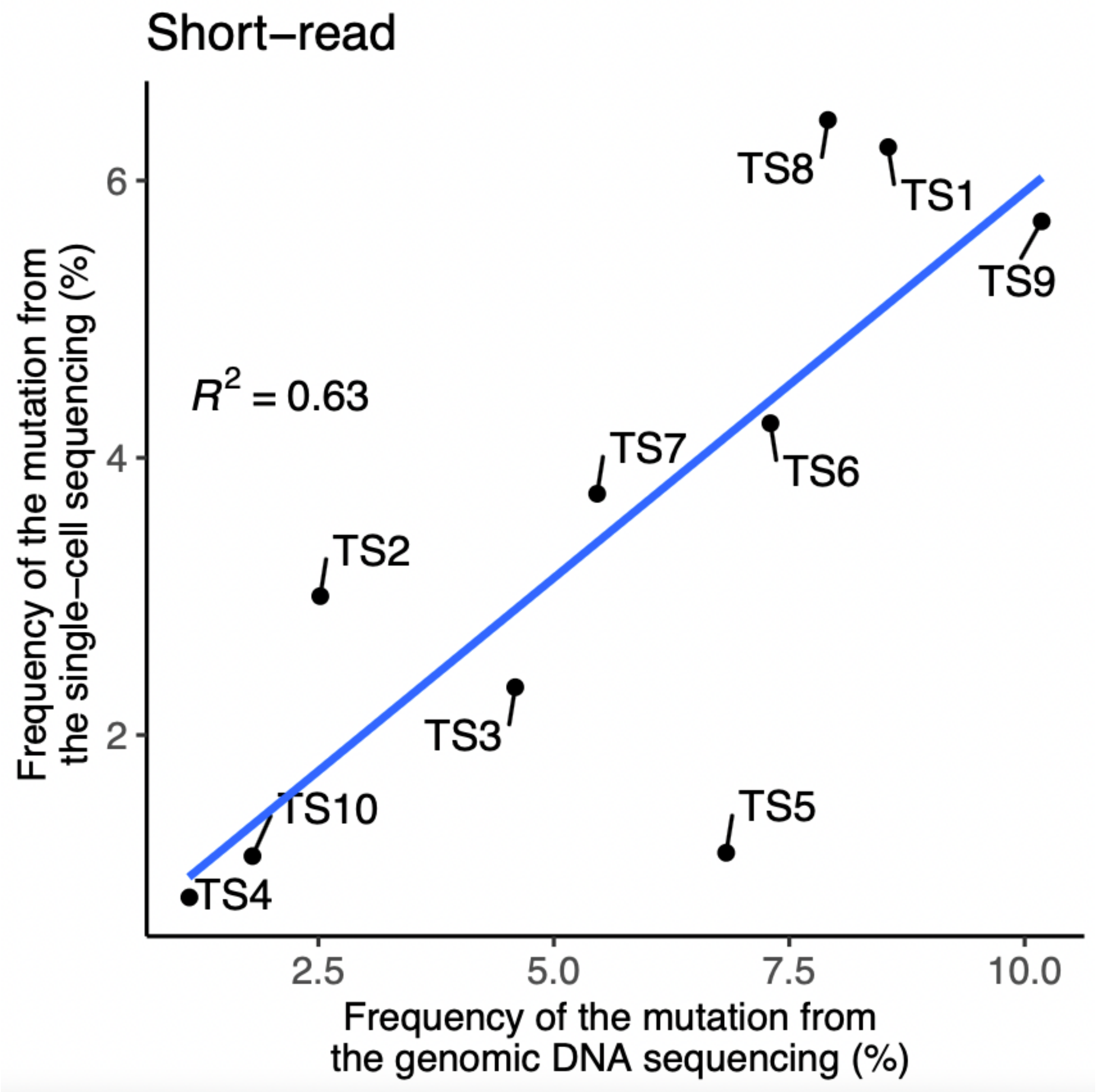

**Supplementary Figure 2. Statistics of sgRNA libraries targeting *TP53* genetic variants.**

| | sgRNA | Targetable variants | Possible variants |
|---|---|---|---|
| NGG_Base_Editors | 74 | 99/351(28.21%) | 920 |
| NG_Base_Editors | 88 | 159/351(45.30%) | 1999 |
| Total | 162 | 251/351(71.51%) | 2892 |

**Supplementary Figure 3. Multiple genetic variants can be introduced by one sgRNA.** Underlines indicate each triplet codon and number indicate position of the codon. Red DNA sequences indicate substituted bases and blues indicate PAM sequences.

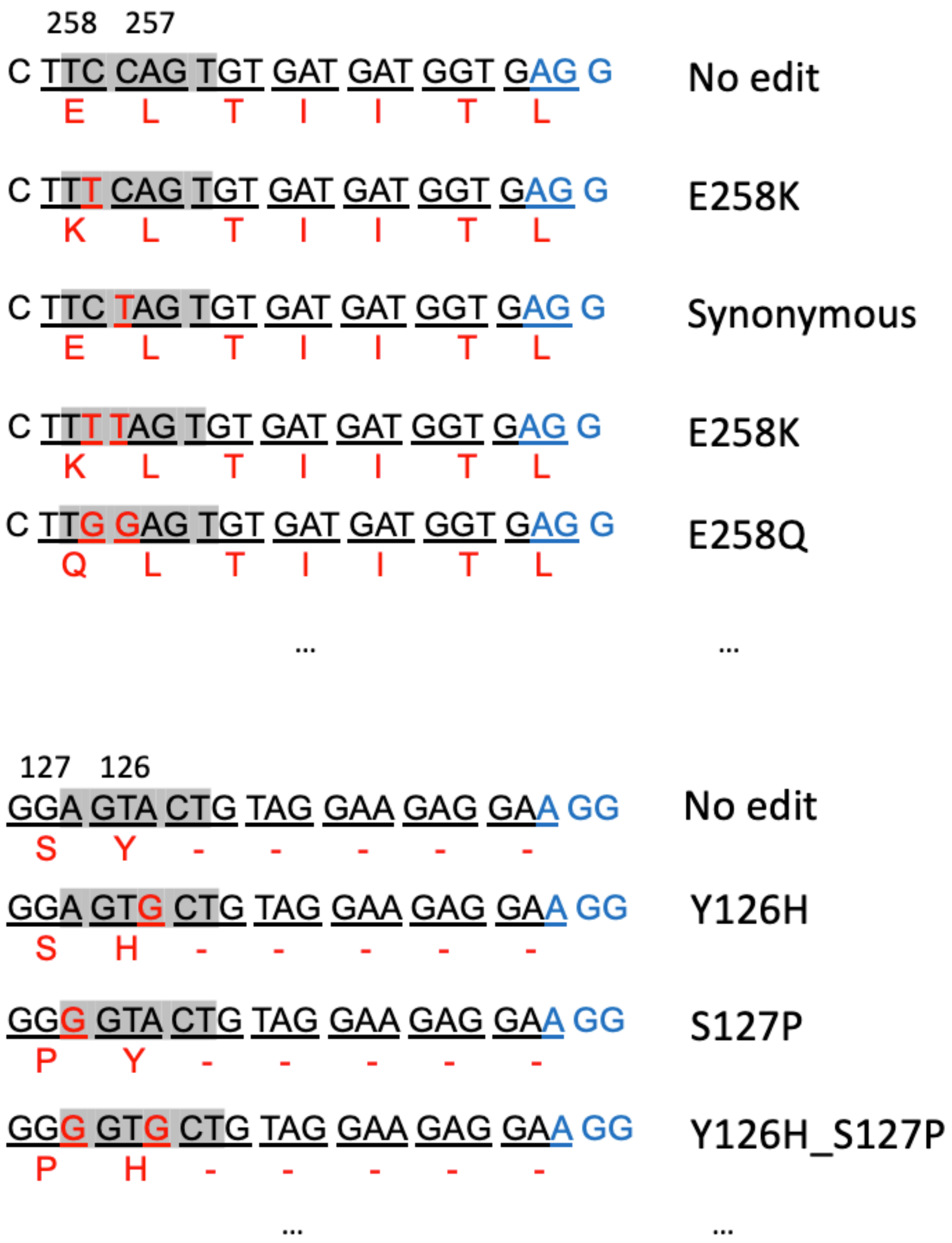

**Supplementary Figure 4. Dot plot showing the proportion of each genetic variant detected from single-cell cDNA and genomic DNA.** Genetic variants generating premature stop codon are removed.

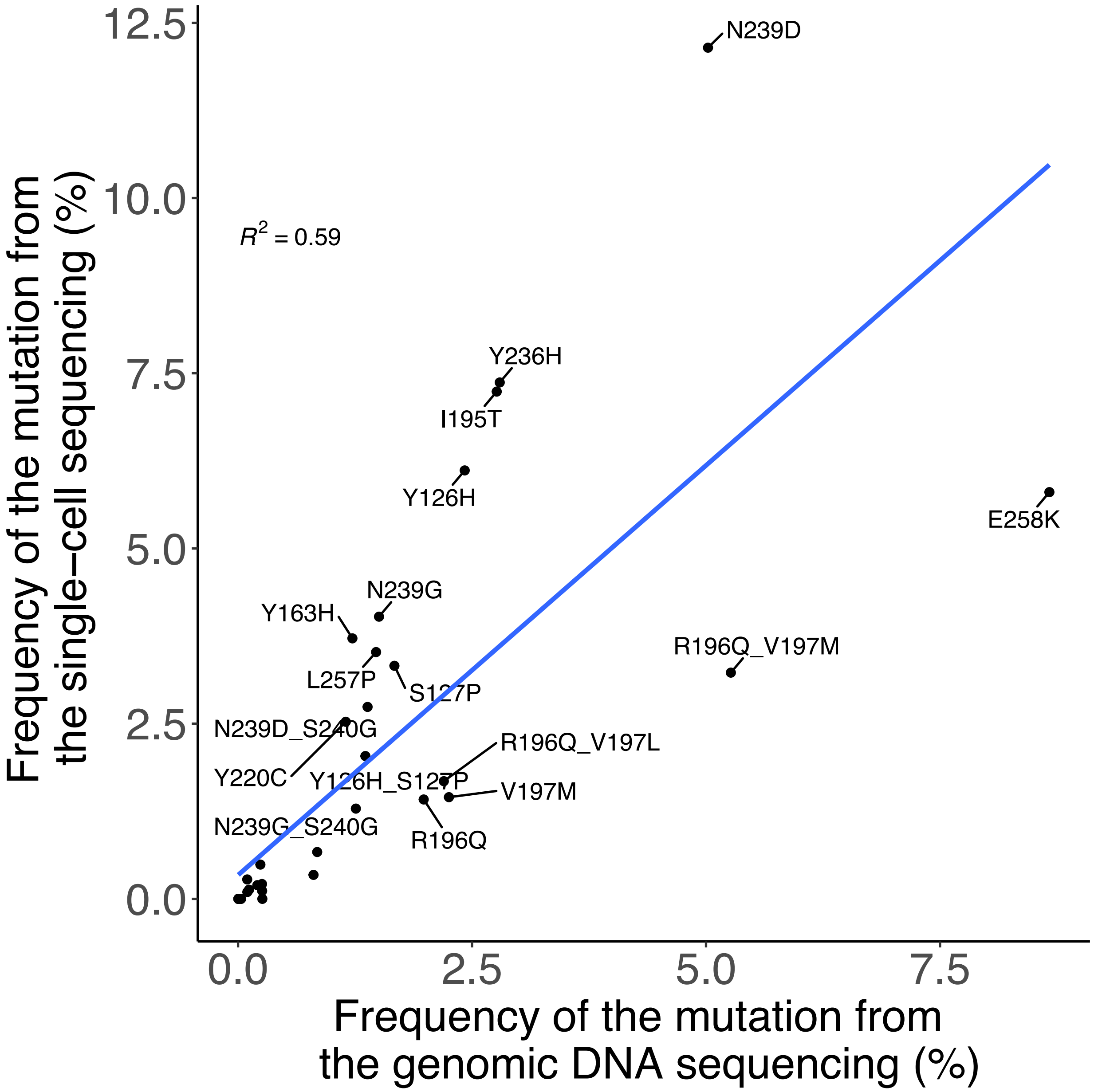

**Supplementary Figure 5. Pie charts showing the proportion of resultant amino acid changes from cells with sgRNA targeting N239D or S127P mutations.** Proportions of mutations are calculated from the single-cell cDNA long-read sequencing. Underlines indicate each triplet codon and number indicate position of the codon. Red DNA sequences indicate substituted bases and blues indicate PAM sequences.

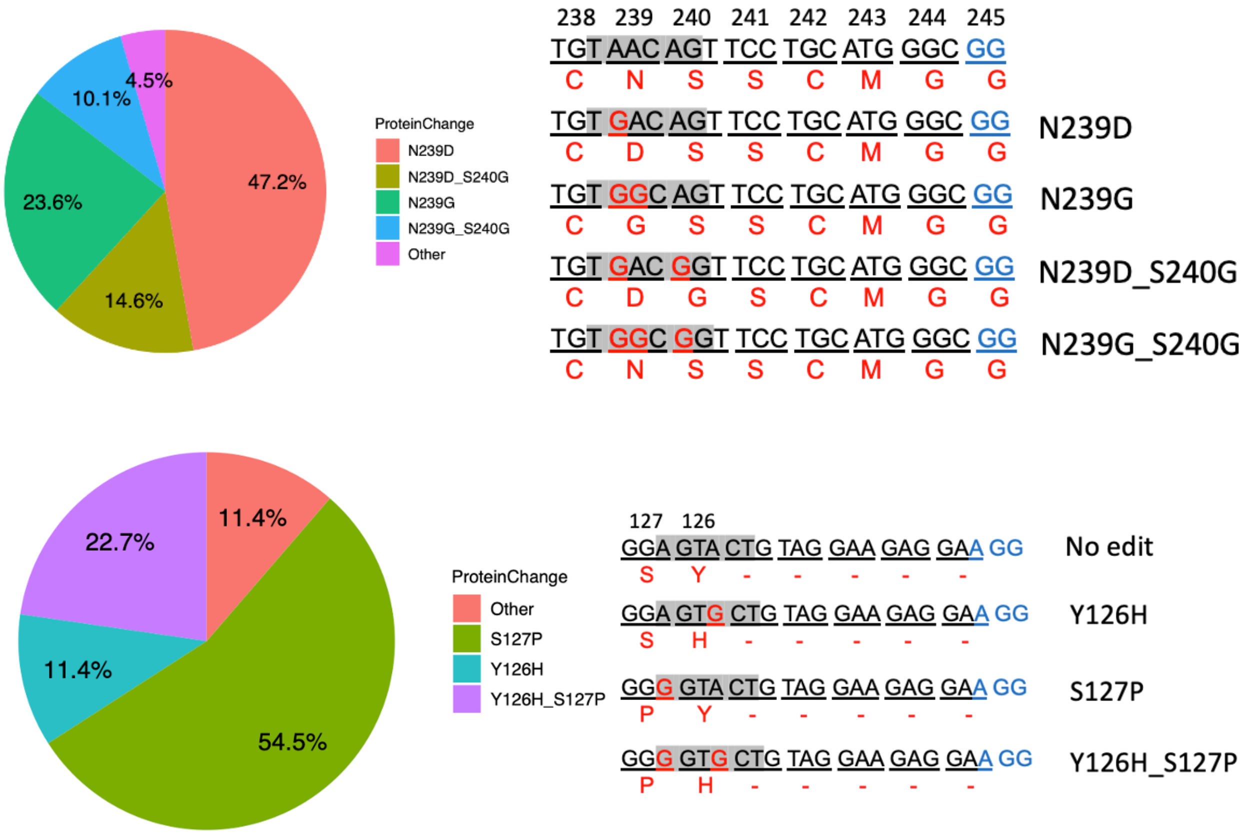

**Supplementary Figure 6. UMAP embedding of cells colored by P53 pathway gene scores.** *TP53* variants were introduced to HCT116 cells using a subset of our sgRNA library and analyzed.

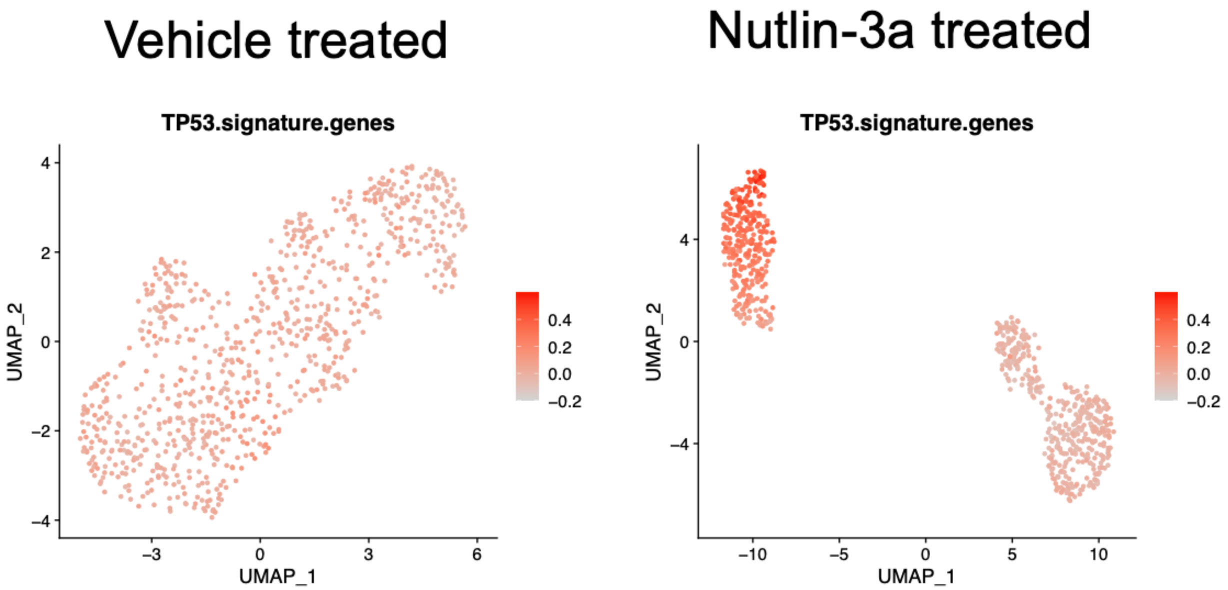

**Supplementary Figure 7. UMAP plot of HCT116 cells which various *TP53* genetic variants are introduced by full sgRNA library.**

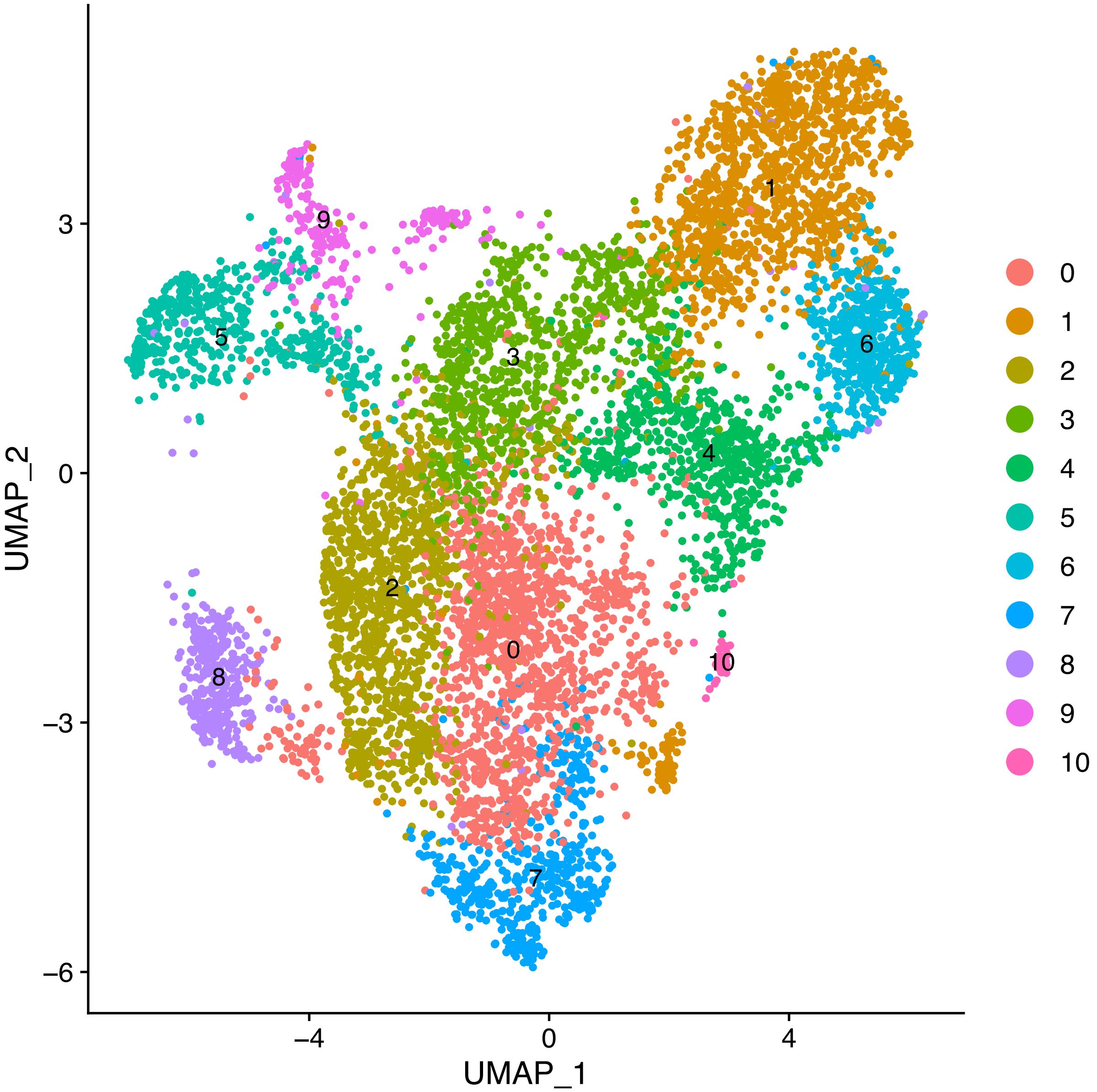

**Supplementary Figure 8.** ***CDKN1A* expression level from HCT116 cells with various TP53 genetic variants.** (A) UMAP embedding of cells colored by *CDKN1A* gene expression. (B) Violin plot showing *CDKN1A* gene expression level per cells with each genetic variant. Reds indicate wild type like variants.

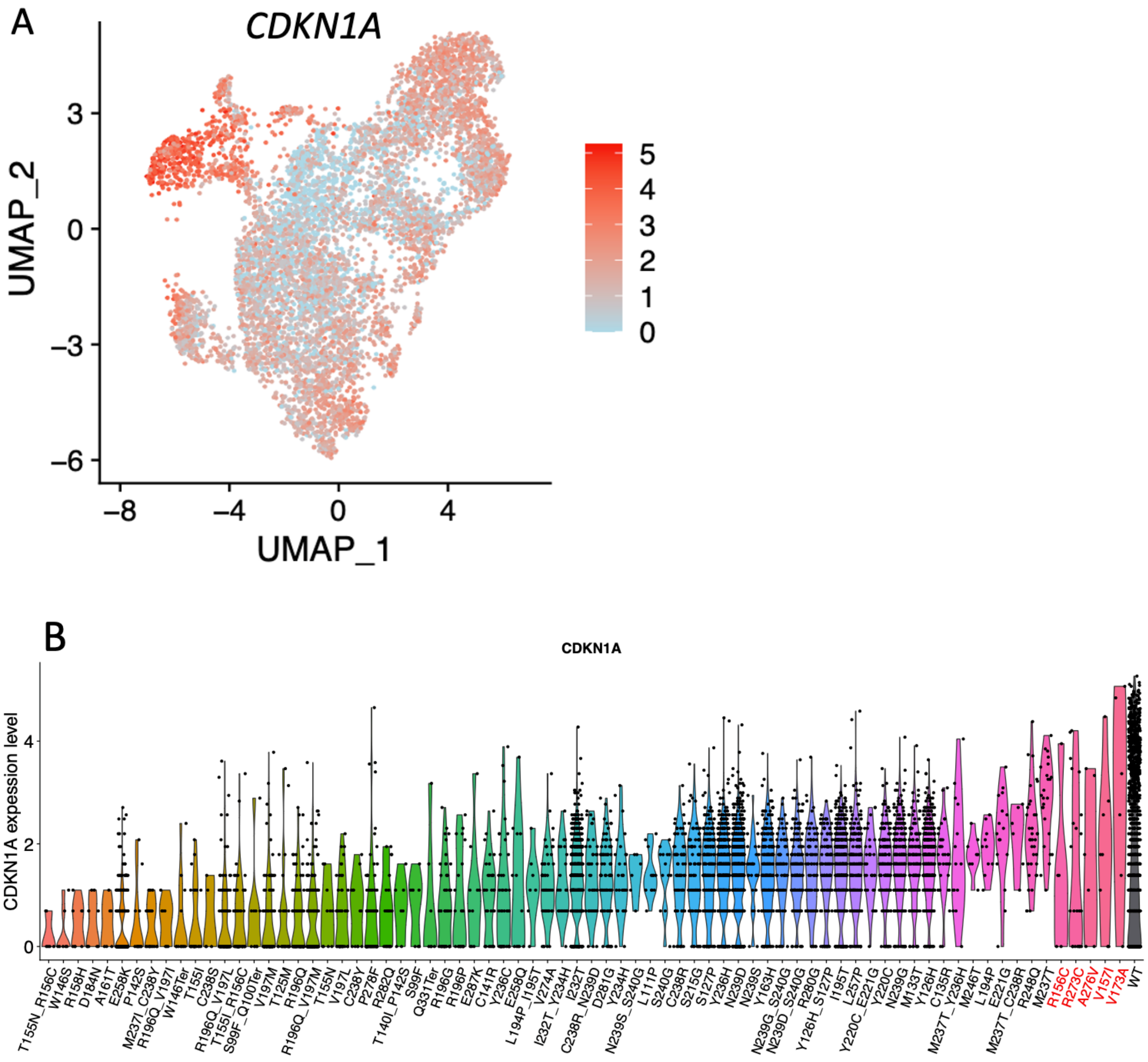

**Supplementary Figure 9. UMAP plot of U2OS cells which various *TP53* genetic variants are introduced by full sgRNA library.**

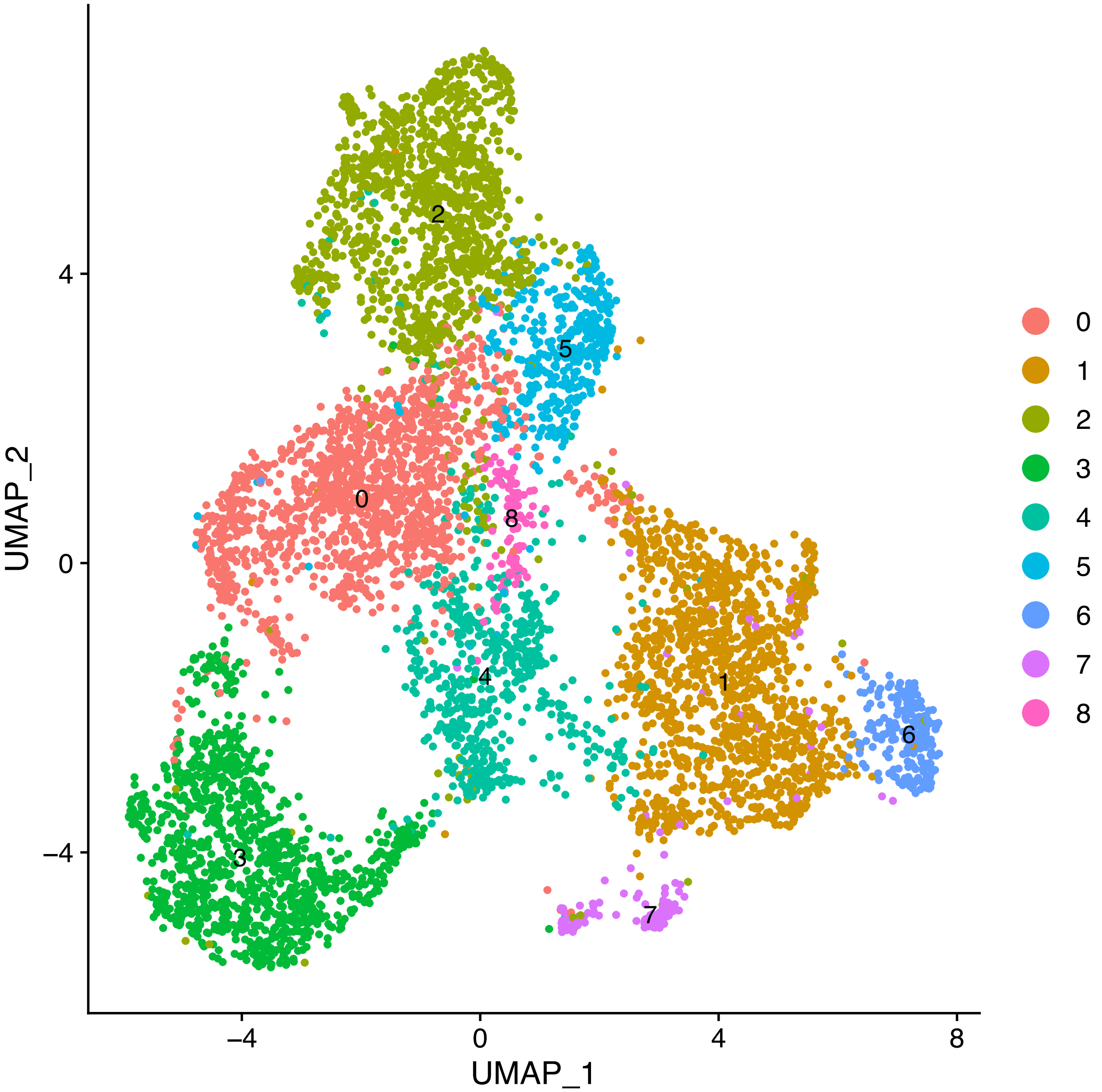

**Supplementary Figure 10.** ***CDKN1A* expression level from U2OS cells with various TP53 genetic variants.** (A) UMAP embedding of cells colored by *CDKN1A* gene expression. (B) Violin plot showing *CDKN1A* gene expression level per cells with each genetic variant. Reds indicate wild type like variants.

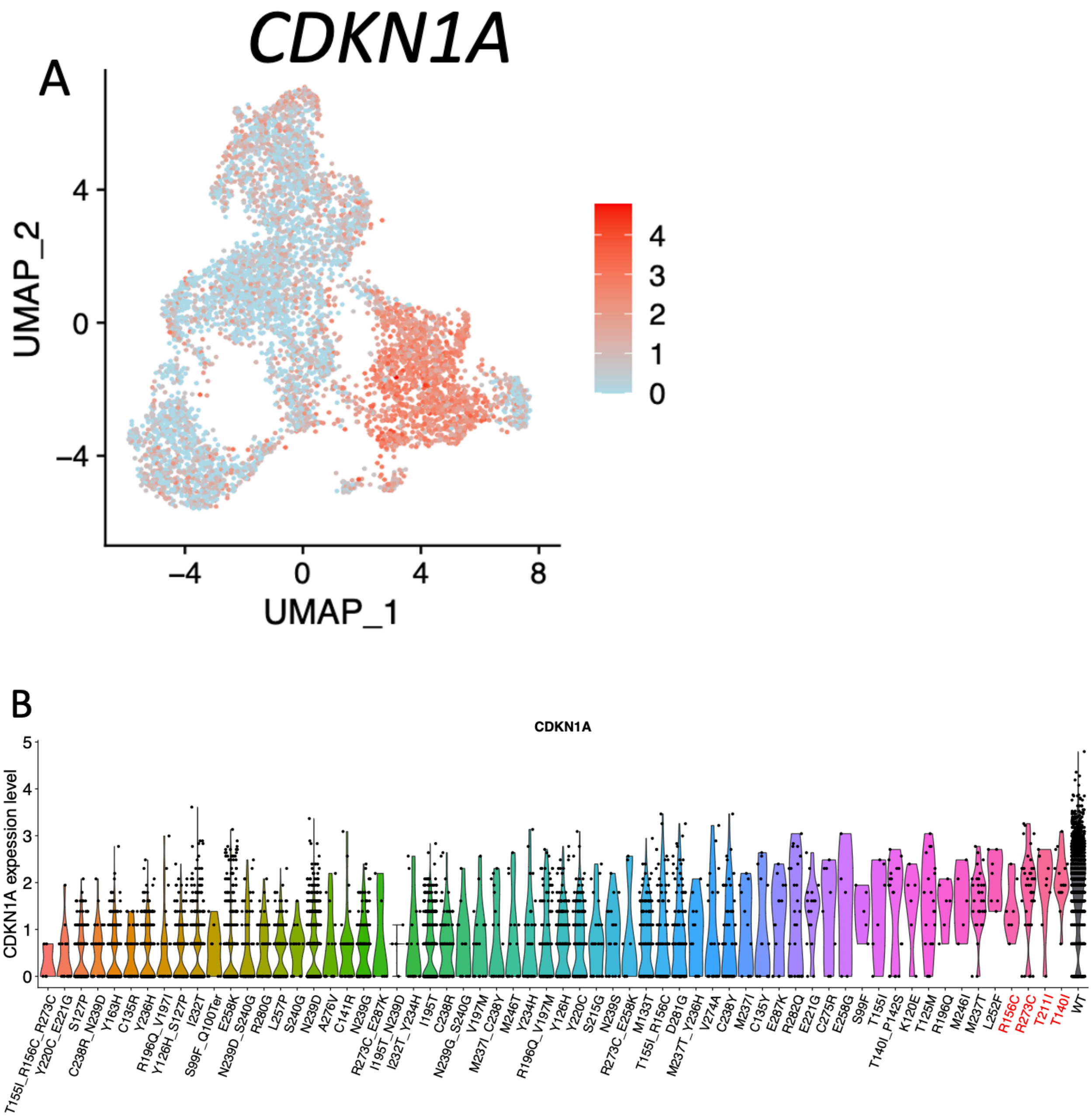

**Supplementary Figure 11. Confirmation of single-cell variants screen using clonal cell-lines.** (A) Overview of the confirmation experiments. (B) Violin plot showing the expression of *CDKN1A* from single-cell sequencing result. (C) Identified genetic variants from bulk RNA-seq from isolated clonal cells. (D) Representative FACS result images showing cell cycle differences in clonal cells.

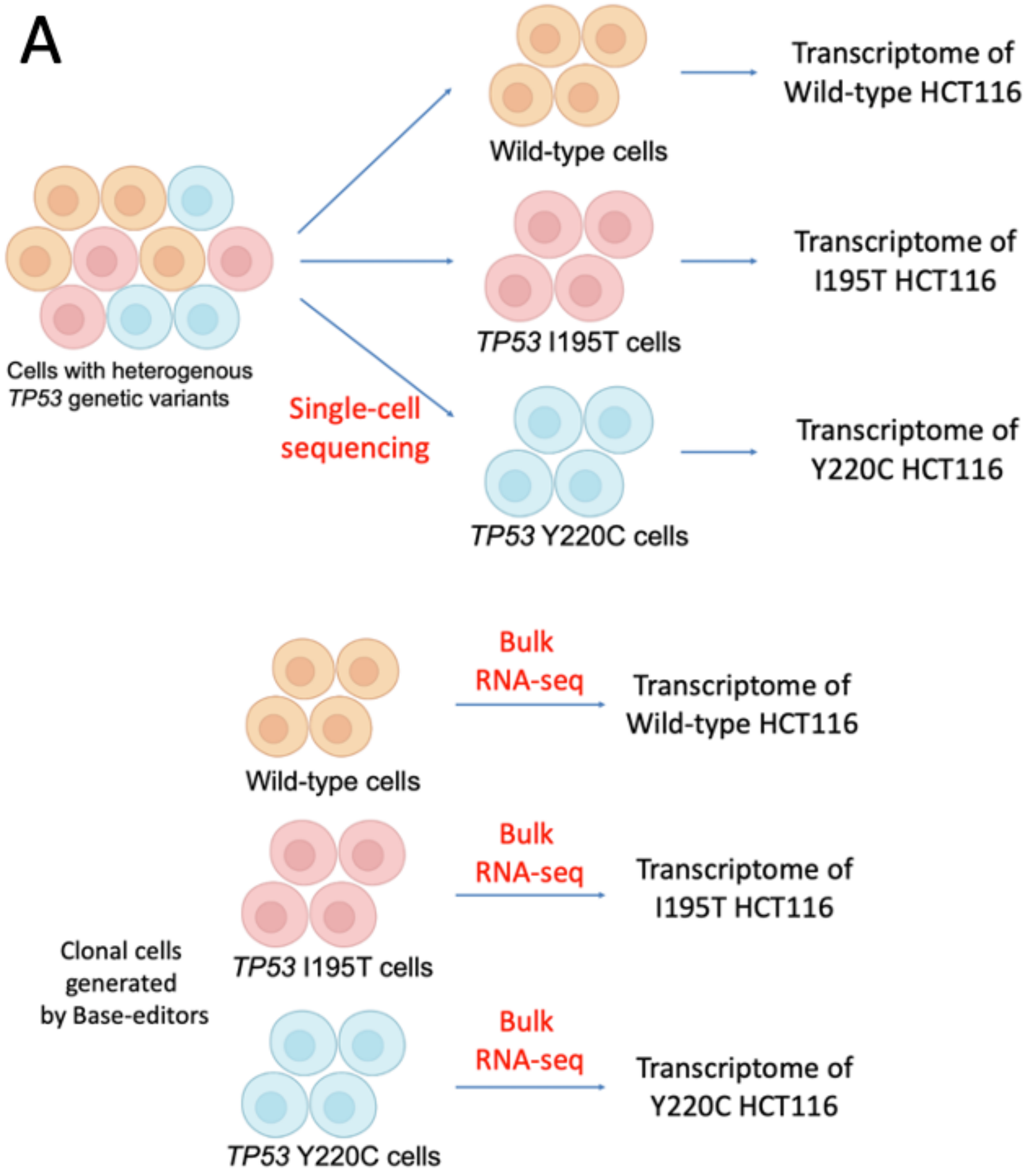

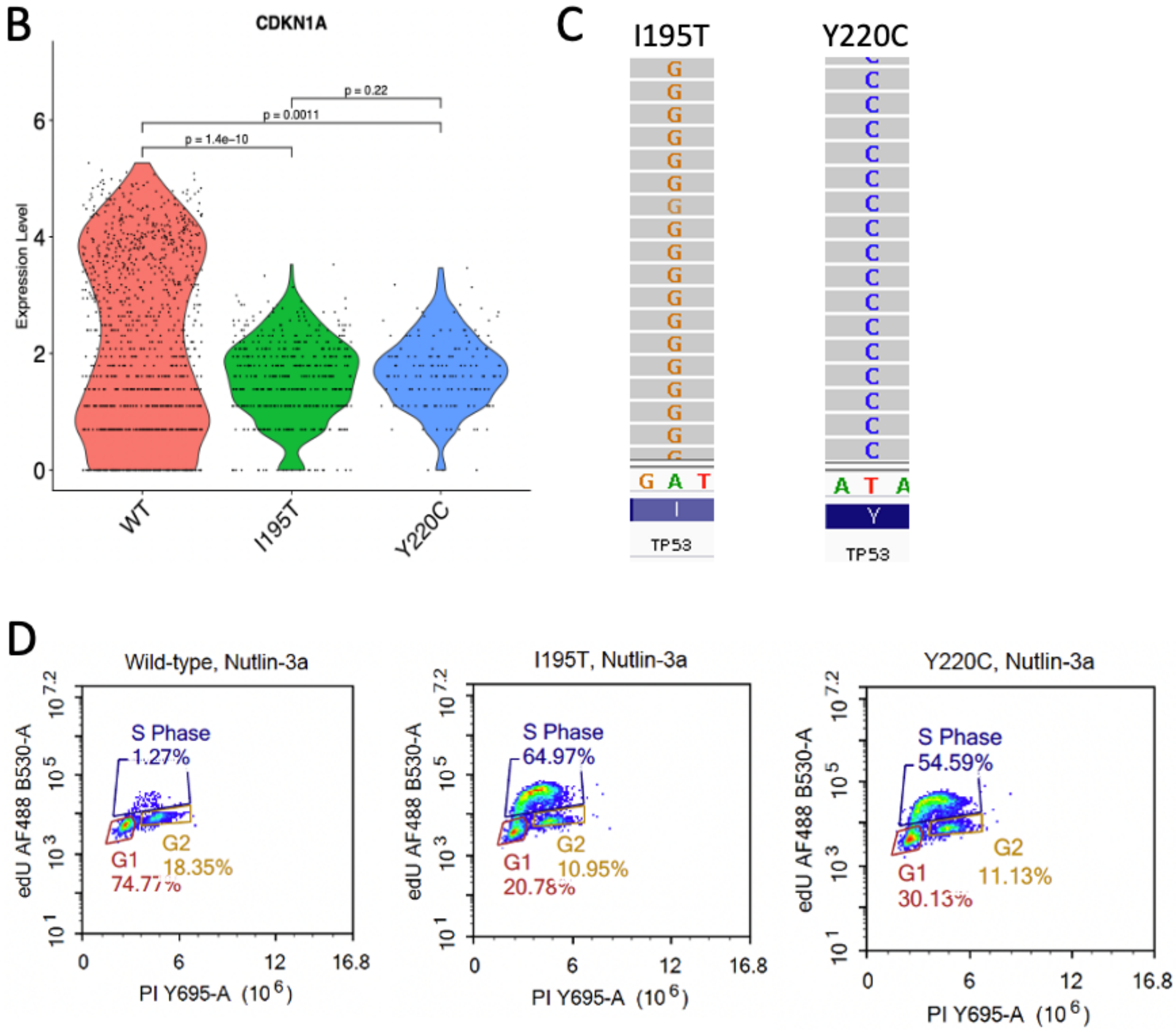
B
CDKN1A
p = 0.22
p = 0.0011
p = 1.4e−10
Expression Level
WT
I195T
Y220C
C
I195T
Y220C
G A T
I
TP53
A T A
Y
TP53
D
Wild-type, Nutlin-3a
I195T, Nutlin-3a
Y220C, Nutlin-3a
S Phase
1.27%
64.97%
54.59%
G1
74.77%
20.78%
30.13%
G2
18.35%
10.95%
11.13%
edU AF488 B530-A
PI Y695-A (10⁶)
